# Genome-wide associations of leaf spectral variation in MAGIC lines of *Nicotiana attenuata*

**DOI:** 10.1101/2023.10.03.560760

**Authors:** Cheng Li, Ewa A. Czyż, Bernhard Schmid, Rishav Ray, Rayko Halitschke, Ian T. Baldwin, Michael E. Schaepman, Meredith C. Schuman

**Affiliations:** Department of Geography, University of Zürich, Winterthurerstrasse 190, 8057 Zürich, Switzerland; Department of Molecular Ecology, Max Planck Institute for Chemical Ecology, Hans-Knoell-Strasse 8, 07745 Jena, Germany; Department of Chemistry, University of Zürich, Winterthurerstrasse 190, 8057 Zürich, Switzerland

**Keywords:** Field spectroscopy, Genome-wide association studies (GWAS), Hierarchical Spectral Clustering with Parallel Analysis, Leaf reflectance, Multiparent Advanced Generation Inter-Cross (MAGIC), *Nicotiana attenuata*

## Abstract

The application of in-field and aerial spectroscopy to assess functional and phylogenetic variation in plants has led to novel ecological insights and supports global assessments of plant biodiversity. Understanding how plant genetic variation influences reflectance spectra will help harness this potential for biodiversity monitoring and improve understanding of why plants differ in functional responses to environmental change. Here, we use a well-resolved genetic mapping population derived from Multi-parent Advanced Generation Inter-cross (MAGIC) lines of *Nicotiana attenuata* to associate genetic differences with differences in leaf spectra between plants in a field experiment in their natural environment. We analyzed the leaf reflectance spectra using a hand-held spectroradiometer (350–2500 nm) on 616 fully genotyped plants of *N. attenuata* grown in a randomized block design. We tested three approaches to conducting Genome-Wide Association Studies on spectral variants. We introduce a new Hierarchical Spectral Clustering with Parallel Analysis (HSC-PA) method. This method efficiently captured the variation in our high-dimensional dataset and allowed us to discover a novel association, between a locus on chromosome 1 and the 734-1143 nm spectral range, spanning the red-edge and near-infrared regions that are sensitive to leaf structure and photosynthetic activity. This locus contains a candidate gene annotated as carbonic anhydrase, an enzyme involved in CO₂ hydration and regulation of photosynthetic efficiency, suggesting a physiological link between variation in leaf optical properties and carbon assimilation. In contrast, an approach treating single wavelengths as phenotypes identified the same associations as HSC-PA, but without the statistical power to pinpoint significant associations. An index-based approach, which reduces complex spectra to a few dimensionless variables, detected two significant associations for *ARDSI*_*C*_w_ (a water-content-related index) with loci on chromosome 1 near genes annotated as a Zeta toxin domain-containing protein, and an Exocyst subunit Exo70 family protein. While these findings are biologically plausible, they represent a very narrow subset of the spectral variation captured by HSC-PA. The HSC-PA approach supports a comprehensive understanding of the genetic determinants of leaf spectral variation which is data-driven but human-interpretable, and lays a robust foundation for future research in linking plant genetics with biodiversity monitoring, large-scale ecological assessment and remote-sensing applications.

## 1 INTRODUCTION

Genetic diversity is essential for the survival and adaptation of species to changing environmental conditions. Radiation reflected, absorbed, and transmitted by plants constitutes the basis for remote sensing of vegetation, commonly in the range from visible to infrared wavelengths corresponding to solar radiation (Knipling 1970; Thenkabail, et al. 2018). Remote sensing may revolutionize the way we study genetic diversity by offering non-invasive methods with the potential of repeated measurement on large spatial scales (Madritch et al. 2014). Remote sensing technologies have not only facilitated the study of Earth’s surface features, including vegetation, water bodies, and land-use patterns, but are also gaining applications in biodiversity research. Notably, high-throughput hyperspectral phenotyping has long been used in crop quantitative genetics to link genetic variation with spectral traits, demonstrating the heritability of multiple biochemical and physiological traits estimated from spectral signatures—thus providing valuable methodological precedents for biodiversity and ecological studies (Grzybowski et al., 2021). Wang and Gamon recently discussed concrete ways in which remote sensing makes unique contributions to monitoring plant biodiversity (Wang and Gamon 2019).

For example, Féret and Asner showcased the capabilities of high-fidelity imaging spectroscopy for mapping the diversity of tropical forest canopies (Asner et al., 2014). Although they do not capture the structural variation of whole canopies, reflectance spectra from single leaves, which can be captured with controlled lighting and background from plants in field conditions, also offer insights into plant physiology, stress responses, and genetic variation, with recent studies establishing a correlation between genetic diversity and leaf reflectance spectra (Serbin et al., 2014; Cavender-Bares et al., 2016; Wang et al., 2018; Asner et al., 2015; Stasinski et al., 2021; Czyż et al., 2023). We recently showed that genetically variable plant populations are also spectrally more variable in comparison to replicates of an inbred genotype grown under either glasshouse or field conditions, while isogenic plants differing primarily in their gene expression are spectrally similar to replicates of the inbred genotype from which they were derived (Li et al., 2023).

A Genome-Wide Association Study (GWAS) is a powerful tool used to identify genetic variants associated with specific traits or diseases in populations. By examining the entire genome, researchers can pinpoint specific genetic markers that correlate with phenotypic variation. A primary advantage of GWAS is that it requires no prior knowledge of potential candidate genes and can be used for the discovery of novel genetic associations (Visscher et al., 2017; Santure et al., 2018). The corresponding disadvantage of GWAS is that it does not provide causal links and thus the mechanisms underlying statistical associations must be dissected and tested. A primary concern of GWAS is the influence of population structure, which, if not addressed, may produce misleading associations (Price et al., 2006). Population structure refers to the presence of subgroups within a population that differ in allele frequencies due to shared ancestry. In GWAS, population structure can be confounding, because genetic variants associated with the subgroup, rather than the trait of interest, may result in false-positive associations. To mitigate this, various methods, such as principal component analysis (PCA) and mixed linear models (Zhang et al., 2010), are employed to correct for population structure in GWAS analyses. Phenotyping remains a substantial bottleneck in genetic studies due to its labor-intensive and time-consuming nature. High-throughput phenotyping, especially hyperspectral imaging, offers a powerful solution— enabling rapid, scalable, and non-invasive measurements of plant traits that can accelerate genotype–phenotype association studies (Furbank & Tester, 2011; Zhang & Zhang, 2018).

*Nicotiana attenuata* Torr. ex S. Watson is a model wild plant for molecular ecology native to the Great Basin Desert of the southwestern USA. It is primarily found in large ephemeral populations following fires in sagebrush and pinyon-juniper ecosystems. Its unique germination behavior is stimulated by cues found in wood smoke and the removal of inhibitors from unburned litter, allowing it to thrive in post-fire environments (Bahulikar et al., 2004).

Ecologically, *N. attenuata* is a compelling subject for its intricate defense mechanisms against herbivores, its tissue-specific diurnal metabolic rhythms, and its ability to acclimate to varying environmental conditions, including high UVB radiation (Glawe et al., 2003; Kim et al., 2011; Li et al., 2016; INH, Galis, and Baldwin 2013). Further, it offers unique opportunities for genetic research due to a well-defined MAGIC (Multiparent Advanced Generation Inter-Cross) population (Ray et al., 2019, 2023). This design comprises 26 phenotypically and genetically differentiated parental lines (PLs) intercrossed to produce two replicate sets each of 325 recombinant inbred lines (RILs; 616 genotyped plants of the total of 650 used in the present study), allowing for the study of phenotype-genotype associations without the confounding influence of population structure. More details on plant material, field site, and generation of MAGIC RIL Population can be found in Appendix S1: Supporting Text.

In this study, we explore the genetic diversity of the MAGIC population to identify associations between genetic variants and leaf spectral traits, advancing our understanding of genetic influences on complex spectral characteristics. Our methodology involves three approaches: analyzing spectral indices, single wavelengths (SW), and implementing Hierarchical Spectral Clustering with Parallel Analysis (HSC-PA). Through GWAS using 168,903 SNPs that showed variation within the population, we investigate genetic links with genome annotation. Our methodological approach and the tools utilized are summarized in Figure 1.

**Figure 1.**
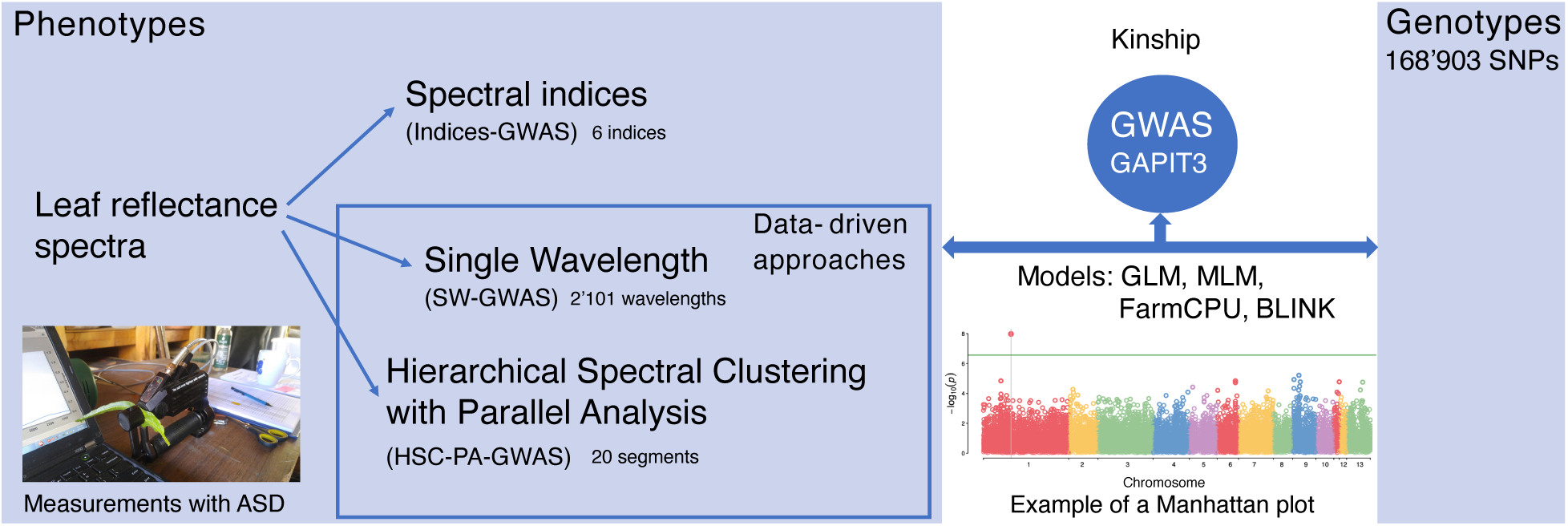
Approaches and toolbox of this study. Photo credit: Meredith C. Schuman.

## 2 MATERIALS AND METHODS

This study’s methodology extends from our previous work (Li et al., 2023), with detailed description available in Appendix S1: Supporting Text. Briefly, our study utilizes two replicate sets each of 325 *N. attenuata* MAGIC RILs, derived from 26 parental lines and six generations of inbreeding, yielding a total of 650 genetically different plants of which 616 were planted in the current experiment. The RILs sequencing was performed at Novogene HK (Ray et al., 2023), and leaf optical properties (350-2500 nm) were measured using a FieldSpec 4 spectroradiometer (Li et al., 2023).

Data analysis was performed using R (R Core Team 2023), employing the spectrolab package (version 0.0.10, Meireles, et al., 2017) and a three-step filtering approach for outlier removal (Li et al., 2023). We employed a linear model to remove known environmental effects from the raw spectral data. Specifically, for each wavelength, we fitted the model:

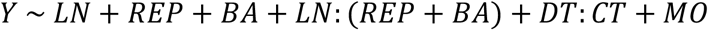

where *LN* represents the influence of the measurement technique (the number of measured leaves), *REP* accounts for potential differences between the two replicate sets of RILs (that occupied different sectors of the field plot and were measured in series), *BA* represents spatial patch effects characterized from *N. attenuata* reference measurements (in each square of 2 × 2 planting positions one was occupied by a “phytometer” plant of a single tester genotype of the species), *LN*: (*REP* + *BA*) models the interaction between measurement technique and spatial effects, *DT*: *CT* captures temporal effects (*DT* for day and *CT* for time of day), and *MO* denotes the maternal cytoplasmic effects. The residuals from each fitted model were taken as the adjusted spectral values, corrected for (i.e., removing) the modeled environmental influences.

We tested three ways of treating the spectral phenotypic data. First, we used six selected spectral indices, which are single values derived from ratios of reflectance at specific wavelengths representing spectral features (details see Table 1 and Appendix S1: Supporting Text). These indices span pigment-related (e.g., chlorophyll a/b, carotenoids), water-related, and structural spectral features across the VIS, red-edge, NIR, and SWIR ranges, thus capturing major axes of leaf functional variation. The indices were first calculated from the raw spectral data and subsequently adjusted using the same linear model as described above. Second, we used single wavelengths (SW), i.e., treating each wavelength in the adjusted spectrum as a phenotype. Third, we used Hierarchical Spectral Clustering with Parallel Analysis (HSC-PA), a new data-driven dimension reduction approach, developed from a method for the analysis of genetic associations with human facial features (Claes et al., 2018; Sero et al., 2019), which accounts for correlations within spectral data while retaining interpretable features. We applied HSC-PA to the adjusted spectral data for downstream analyses. We associated the resulting phenotypic data with 168,903 SNPs using multiple GWAS models based on the Genome Association and Prediction Integrated Tool (GAPIT) version 3 (Wang and Zhang 2021). The GWAS models include: General Linear Model (GLM, Price et al., 2006), Mixed Linear Model (MLM, Yu et al., 2006), Fixed and random model Circulating Probability Unification (FarmCPU, Liu et al., 2016), and Bayesian information and Linkage-disequilibrium Iteratively Nested Keyway (BLINK, Huang et al., 2019). Kinship matrices were automatically calculated by GAPIT using the Zhang method (Zhang et al., 2010) from the genotype data; these matrices were applied only in models that incorporate kinship (i.e., MLM). See more details of the four models in Appendix S1 Supporting Text.

**Table 1.** Six Spectral Indices Used in Indices-GWAS.

| Index | Formula | Example of Validated Systems | Citations |
| --- | --- | --- | --- |
| *NDWI* | $\frac{R_{865} - R_{1614}}{R_{865} + R_{1614}}$ | Crops, forests, wetlands, grasslands | (Gao 1996; F. Zhang and Zhou 2019; Karthikeyan, Chawla, and Mishra 2020; Huete 2012; Adam, Mutanga, and Rugege 2010; Gu et al. 2007; Helfenstein et al., 2022) |
| *CI<sub>re</sub>* | $\frac{R_{783}}{R_{704}} - 1$ | Crops, trees and vines, wetlands | (Gitelson, Merzlyak, and Chivkunova 2001; Qian et al. 2021; Gitelson et al. 2005; Gitelson, Gritz, and Merzlyak 2003; DeLancey et al. 2019; Helfenstein et al., 2022) |
| *CCI* | $\frac{R_{560} - R_{664}}{R_{560} + R_{664}}$ | Crops, trees, forests | (Gamon et al. 2016; Dechant et al. 2020; Grabska et al. 2019; Helfenstein et al., 2022) |
| *ARDSI<sub>C<sub>ab</sub></sub>* | $\frac{R_{750} - R_{730}}{R_{770} + R_{720}}$ | Crops | (Wan et al. 2021) |
| *ARDSI<sub>C<sub>w</sub></sub>* | $\frac{R_{1360} - R_{1080}}{R_{1560} + R_{1240}}$ | Crops | (Wan et al. 2021) |
| *ARDSI<sub>C<sub>m</sub></sub>* | $\frac{R_{2200} - R_{1640}}{R_{2240} + R_{1720}}$ | Crops | (Wan et al. 2021) |

Multiple testing adjustments were conducted on 168,903 SNPs. Initially, an exploratory threshold of *p* < 1 × 10^−5^ was used for preliminary SNP selection. This was followed by a false discovery rate (FDR) adjustment (Benjamini and Hochberg 1995) and the application of the PhenoSpD method (Nyholt 2004; Zheng et al., 2018) to determine the significance threshold (0.05/*M*_*eff*_), taking into account the effective number of independent trait calculations. The PhenoSpD method first derives the pairwise phenotypic correlation matrix from GWAS summary statistics using cross-trait LD Score regression. Specifically, for two traits *t*_1_and *t*_2_, the regression

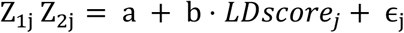

is calculated across SNPs *j*, where *Z*_1j_ and *Z*_2j_ are the z-scores from GWAS summary statistics for each trait, and the intercept *α* provides an estimate of the phenotypic correlation between traits. This phenotypic correlation matrix is then subjected to spectral decomposition (SpD), in which the eigenvalues (*λ*_*i*_) are used to calculate the effective number of independent tests:

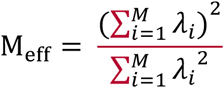

where *M* is the total number of phenotypes. We used 0.05/*M*_*eff*_ as the significant threshold. SNPs with FDR-adjusted p-values below the significance threshold were considered significantly associated. For candidate gene selection, we extracted genes within ±100 kb of significant SNPs from the *N. attenuata* genome, following the approach used in closely related *Nicotiana tabacum* (Lai et al., 2021). We then refined the candidate gene list using the *N. attenuata* Data Hub (http://nadh.ice.mpg.de/NaDH/) to filter out genes lacking transcript accumulation evidence in leaves under various conditions. Genes with relevant functions were then identified based on annotation information.

We also conducted GWAS using indices, single wavelengths, and HSC-PA segments with raw spectral data as a general comparison to assess the effect of removing known environmental effects (see Appendix S1: Supporting Text, Results from raw spectral data).

## 3 RESULTS

### 3.1 Phenotypic analysis

#### 3.1.1 Variance partitioning and adjustment of raw spectra

To advance our understanding of the phenotypic variation in leaf spectral traits and to remove the known environmental effects, we employed linear models. These models partitioned variance across the spectral range of 350–2500 nm, assessing contributions to cumulative multiple *R*^2^values (i.e. percent sum of squares) from measurement technique (*LN*), replicate and spatial effects and their interactions with measurement technique (*REP* + *BA* + *LN*: (*REP* + *BA*)), measurement time (*DT*: *CT*), and maternal cytoplasm (*MO*) within the 616 plants. Figure 2 illustrates these components for each wavelength as stacked bars, which—given the large number of wavelengths—merge into a continuous stacked profile that collectively accounts for the total variance. Analysis of variance results indicated that residual phenotypic variance between plants, representing genetic and residual environmental contributions accounted from 71.7% to 89.5% of the total phenotypic variance.

**Figure 2.**
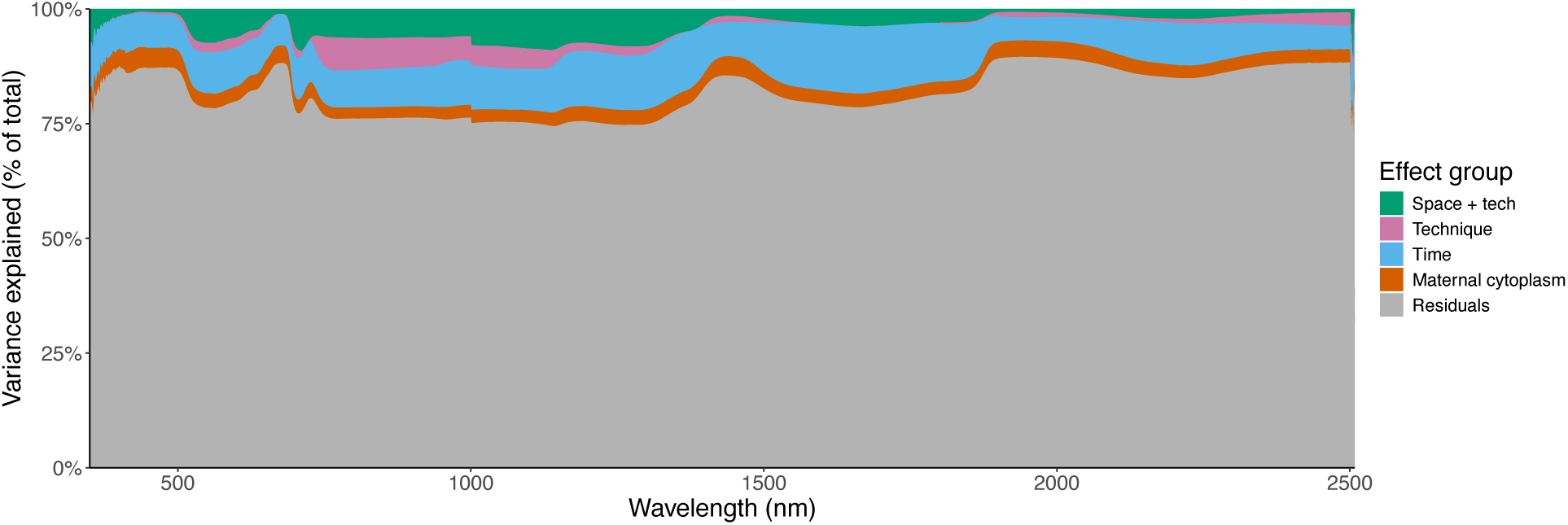
Variance partitioning across the spectral range of 350 to 2500 nm. Each column represents the cumulative percentage of variance for a given wavelength.

#### 3.1.2 Spectral characteristics of leaves

Our analyses focus on calculated leaf reflectance spectra spanning from 400 to 2500 nm, as the initial region (350–399 nm) has a poor signal-to-noise ratio and is commonly excluded from downstream analyses (Cavender-Bares et al., 2016). This range includes the visible (VIS), near infrared (NIR), and shortwave infrared (SWIR) regions. This broad spectral range captured variation among the 616 plants, shown in Figure 3a as an orange shaded region indicating the range of adjusted reflectance spectra for all samples. The blue shaded region indicates the range of raw reflectance spectra before adjustment with linear models. The NIR region showed more consistency in reflectance values among the plants, contrasting with greater variability in the VIS and SWIR regions.

**Figure 3.**
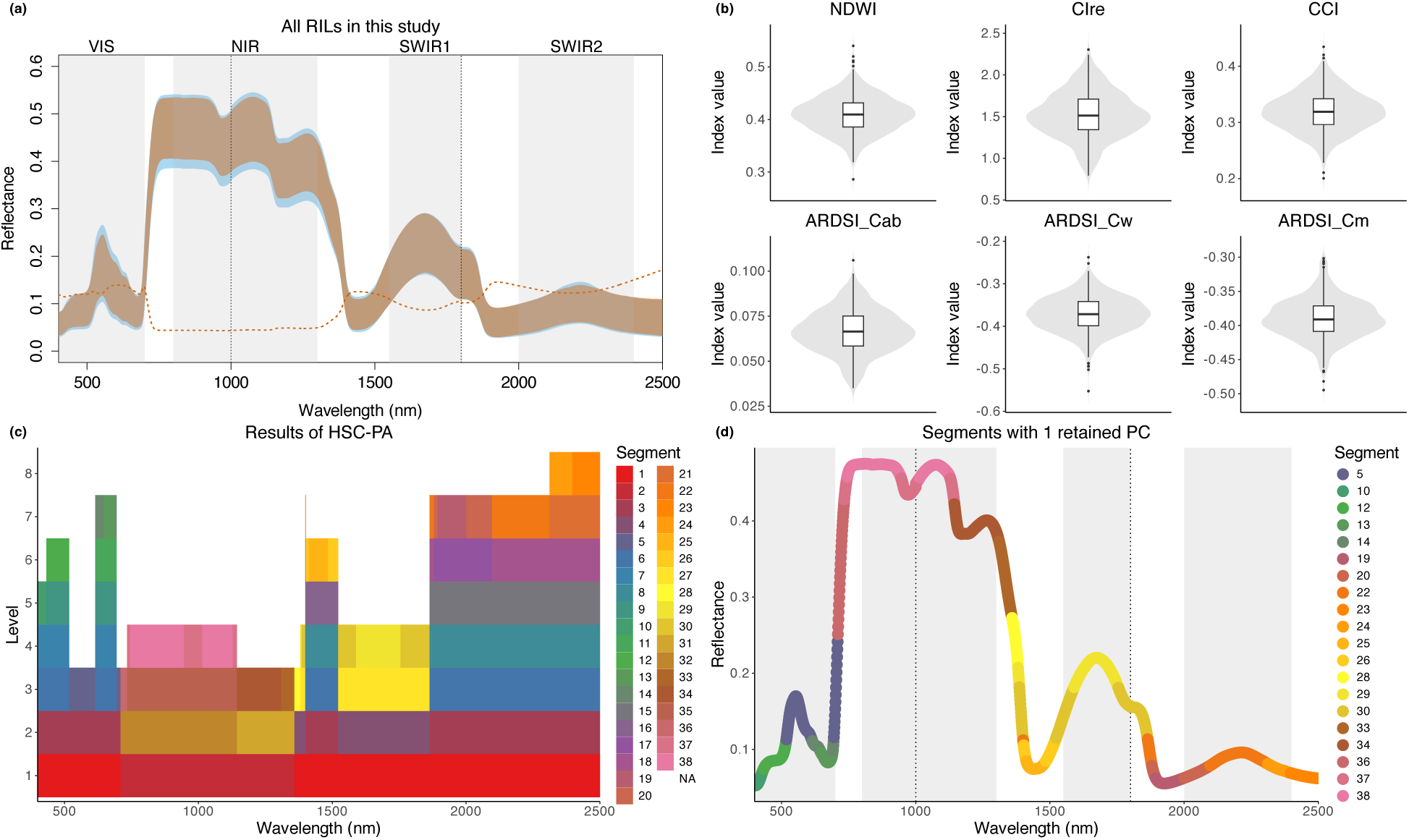
Spectra as phenotypes. (a) Leaf reflectance spectra for all 616 plants examined in this study. The orange shaded region illustrates the range of adjusted spectra, and the blue shaded region the range of raw spectra. The orange dotted line represents the coefficient of variation for the entire sample set. (b) Distribution of the six spectral indices used in the Index-GWAS approach. (c) Results from the Hierarchical Spectral Clustering with Parallel Analysis (HSC-PA). A total of 38 segments, differentiated by color, emerged from the clustering. Regions left blank (NA) indicate segments that retained one Principal Component (PC) in the prior level, halting further clustering. (d) Presentation of the 20 segments, shaped as example spectra, each maintaining one retained PC. Spanning the entire spectrum from 400 to 2500 nm, these segments served as phenotypes in the HSC-PA-GWAS approach to pinpoint associations with genetic variances.

The distribution of six selected spectral indices (Figure 3b), encapsulating attributes like water content and chlorophyll concentration, provided more specific information about leaf physiology. Note that the y-axis represents the index values, which are specific to each index calculation and therefore cannot be directly compared across indices.

Our subsequent application of the HSC-PA segmented the adjusted spectra into 38 distinct patterns. The blank (NA) areas are regions where further clustering was halted due to the retention of a singular Principal Component (PC) from a previous level. This approach allowed for a finer granularity in the spectral data representation while accounting for correlations among wavelengths. The hierarchical clustering process concluded at different levels, ranging from level 3 to level 8, indicating the varying depths of spectral similarities among the segments. Table 2 shows the detailed spectral region and potential associated features of the segments used in HSC-PA-GWAS. Appendix S1: Table S1 shows the spectral regions of all 38 segments.

**Table 2.** Spectral Ranges of Segments Used in GWAS with Potential Associated Features.

| Segment | Spectral range (nm) | Potentially associated physiological features* |
| --- | --- | --- |
| 5 | 518–614, 695–709 | VIS–NIR: Chlorophyll a, chlorophyll b, carotenoids, anthocyanins; leaf structure |
| 10 | 400–432 | VIS: Chlorophyll a, chlorophyll b, carotenoids, anthocyanins |
| 12 | 433–517 | VIS: Chlorophyll a, chlorophyll b, carotenoids, anthocyanins |
| 13 | 648–690 | VIS: Chlorophyll a, chlorophyll b, carotenoids, anthocyanins |
| 14 | 615–647, 691–694 | VIS: Chlorophyll a, chlorophyll b, carotenoids, anthocyanins |
| 19 | 1894–2000 | SWIR: Water content |
| 20 | 1880–1893, 2001–2096 | SWIR: Non-pigment organic composition (e.g., lignin, cellulose, starch, protein, sugar, oil) |
| 22 | 1400–1401, 1864–1879, 2097–2311 | NIR–SWIR: Water content; leaf structure; non-pigment organic composition |
| 23 | 2397–2500 | SWIR: Water content |
| 24 | 2312–2396 | SWIR: Non-pigment organic composition |
| 25 | 1412–1484 | SWIR: Water content |
| 26 | 1402–1411, 1485–1522 | SWIR: Non-pigment organic composition |
| 28 | 1359–1380 | SWIR: Water content |
| 29 | 1381–1385, 1590–1754 | SWIR: Non-pigment organic composition |
| 30 | 1386–1399, 1523–1589, 1755–1863 | SWIR: Non-pigment organic composition |
| 33 | 1311–1358 | NIR: Water content |
| 34 | 1144–1310 | NIR: Leaf structure |
| 36 | 710–733 | NIR: Leaf structure |
| 37 | 734–744, 946–1014, 1129–1143 | NIR: Leaf structure |
| 38 | 745–945, 1015–1128 | NIR: Leaf structure |

Note: The associated features are based on common findings (Jacquemoud and Ustin 2019) and are intended to indicate physiological properties of leaves known to affect the given spectral ranges. These interpretations are subject to various factors and should not be considered definitive.

By visualizing the 20 segments that each retained only a single PC, we were able to span the entire spectrum from 400 to 2500 nm. These segments, parsed into several interpretable regions, were then used for HSC-PA-GWAS approach, offering phenotypic data for uncovering associations with underlying genetic variations.

The distributions of the six indices derived from raw spectra, along with the HSC-PA results based on raw spectral data, are shown in Appendix S1: Figure S1.

#### 3.1.3 Effective number of independent traits (*M*_*eff*_)

To account for multiple testing due to the analysis of multiple phenotypes in our GWAS approaches, we determined the effective number of independent traits, denoted as *M*_*eff*_, using the PhenoSpD method (Zheng et al., 2018). For the Index-GWAS approach, which employed six distinct spectral indices, the effective number of independent tests, *M*_*effiind*_, was determined to be 5.0. In the Single Wavelength GWAS (SW-GWAS) approach, the effective number, *M*_*effsw*_, was notably higher at 999.9. Lastly, for the Hierarchical Spectral Clustering with Parallel Analysis GWAS (HSC-PA-GWAS) approach, which clusters the spectra based on similarity, the effective number, *M*_*effhsc-pa*_, was found to be 10.5, reflecting the reduced dimensionality and the focus on major patterns of variation. These calculated *M*_*eff*_ values are used to set a threshold (0.05/*M*_*eff*_) for significant associations.

### 3.2 SNP marker analysis

The kinship relationships among the 616 plants are visualized in Appendix S1: Figure S2. Through a heatmap, the genetic relatedness between individuals is presented. Most values in the histogram gravitate towards zero, indicating a consistent and minimal genetic difference between plants. This uniform distribution of kinship values affirms the thorough genetic mixing of the parental lines in the MAGIC population and is consistent with the finding that this population does not display obvious structure (Ray et al., 2023).

Appendix S1: Figure S3 presents the frequency of heterozygosity of individual and markers, while Appendix S1: Figure S4 illustrates the linkage disequilibrium (LD) decay over distance.

### 3.3 Results of genome-side association studies (GWAS)

#### 3.3.1 Index-GWAS results

The Manhattan plot presented in Figure 4a, shows the associations between the SNPs and the six indices (*NDWI*, *CCI*, *CI*_*re*_, *ARDSI*_*C*_*αb*_, *ARDSI*_*C*_w_, and *ARDSI*_*C*_*m*_) using four models (GLM, MLM, FarmCPU, and BLINK). Several genomic regions displayed distinct associations according to the −log_10_(p-values). An exploratory threshold, defined at 1 × 10^−5^ (denoted by the grey horizontal line), was adopted to shortlist SNPs for a detailed examination.

**Figure 4.**
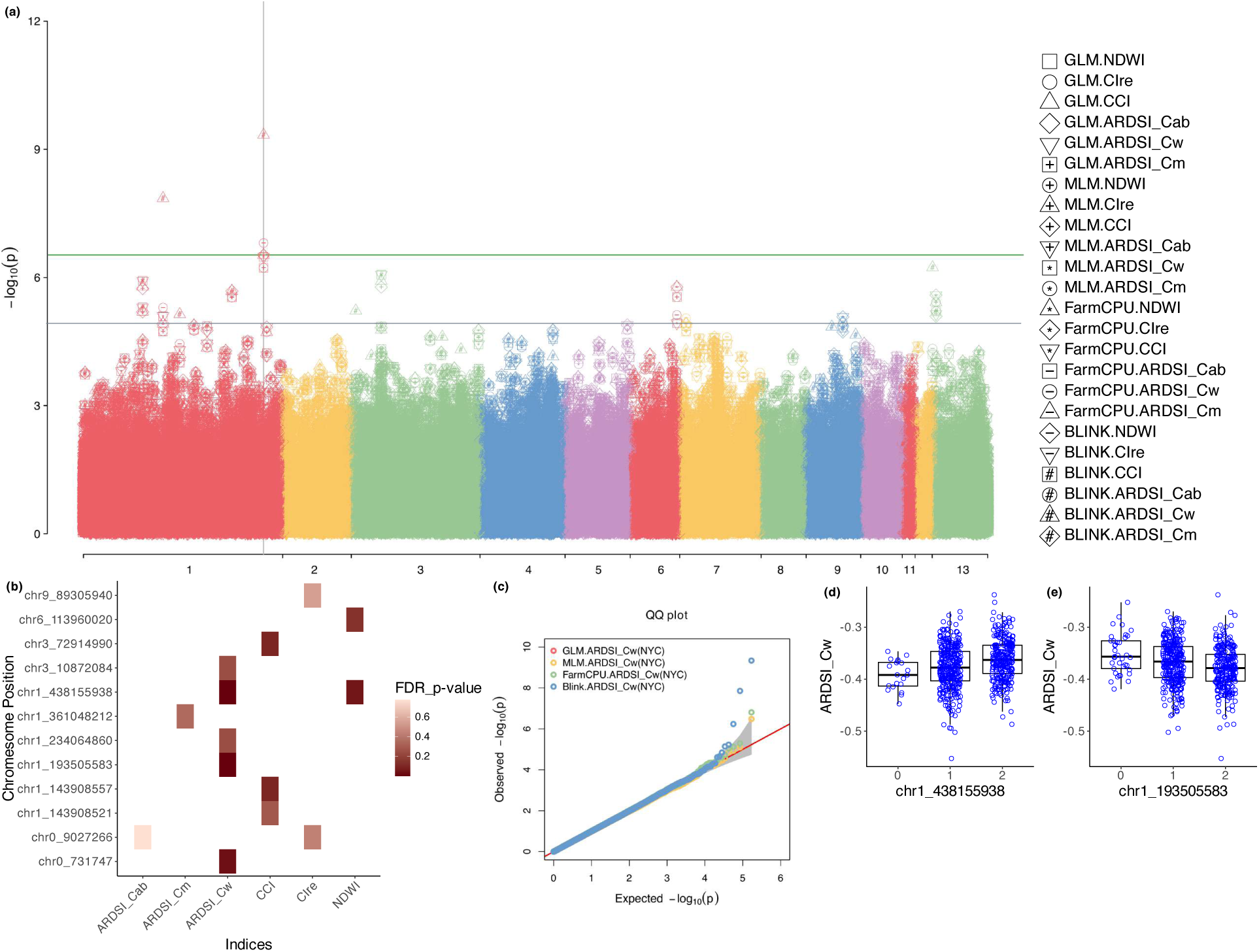
Index-GWAS results. (a) Manhattan plot showing the −*log*_10_(*p*_*value*) of SNP associations for the six indices across various models. The grey horizontal line indicates a p-value of 1 × 10^−5^, which is used to select associations to generate figure (b). (b) Visualization of associations that have p-values less than 1 × 10^−5^. The x-axis represents the six indices, the y-axis the corresponding associated SNPs. The gradient red color scale represents the FDR_adjusted p-values. (c) QQ plot contrasting the observed versus the expected −*log*_10_(*p*_*value*) for the GWAS analysis on the *ARDSI*_*C*_w_ index. The diagonal line represents the expected distribution under the null hypothesis. The different color schemes depict the models used: red for GLM, yellow for MLM, green for FarmCPU, and blue for BLINK. (d)-(e) Phenotypic distribution for two significant markers, chr1_438155938(d) or chr1_193505583(e), in association with *ARDSI*_*C*_w_. 0 represents a homozygous allele from the reference, 1 a heterozygous allele, and 2 a homozygous alternative allele.

In Figure 4b, each of the 6 indices is plotted against the SNPs that exceeded the exploratory threshold. The gradient color scale indicates the association’s intensity, with deeper hues indicating smaller FDR-adjusted p-values. This representation provides a concise overview of the genomic regions that are most strongly correlated with each spectral index. In total, 12 SNPs exhibited p-values less than 1 × 10^−5^: five with *ARDSI*_*C*_w_, three with *CCI*, two each with *NDWI* and *CI*_*re*_ and one each with *ARDSI*_*C*_*m*_and *ARDSI*_*C*_*αb*_. The index *ARDSI*_*C*_w_ displayed two associations lower than the significance threshold: 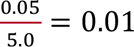, with two SNPs in chromosome 1. The first SNP was located at chr1_193505583, near Niat3g_08138, annotated as a Zeta toxin domain-containing protein. The second was at chr1_438155938, close to Niat3g_17277 (TPR_REGION domain-containing protein) and Niat3g_17282 (Exocyst subunit Exo70 family protein). Furthermore, *ARDSI*_*C*_w_ was chosen as a representative example for the QQ plots in Figure 4c and 4d. This plot serves as a diagnostic tool for the GWAS results. Each model is represented by a unique color: red for GLM, yellow for MLM, green for FarmCPU, and blue for BLINK. The alignment of the observed points with the diagonal line indicates that the GWAS results are mostly consistent with the expectations under the null hypothesis. However, deviations can pinpoint regions with stronger genetic signals. Figure 4e and 4f show the phenotypic distributions for two significant markers, where 0 represents a homozygous reference allele, 1 a heterozygous allele, and 2 a homozygous alternative allele.

#### 3.3.2 SW-GWAS and HSC-PA-GWAS results

In the SW-GWAS analysis, the BLINK model was selected to investigate the genetic correlations with 2101 spectral wavelengths spanning from 400 to 2500 nm. Figure 5a highlights the genetic associations for the spectral wavelength at 952 nm, which displayed the strongest association among all phenotypes, serving as an example. The diagnostic QQ plot for this wavelength is also presented in Figure 5c. Figure 5b offers an in-depth depiction of associations that exceeded the exploratory threshold. The X-axis represents the spectral wavelengths, the Y-axis the associated SNPs, providing a holistic view of the spectral regions correlated with each SNP. A total of 27 SNPs are displayed, with the majority linked to one or multiple continuous spectral regions. This overlap indicates the high correlation inherent in the leaf reflectance spectra data, underscoring the need for methods that can account for these correlations.

**Figure 5.**
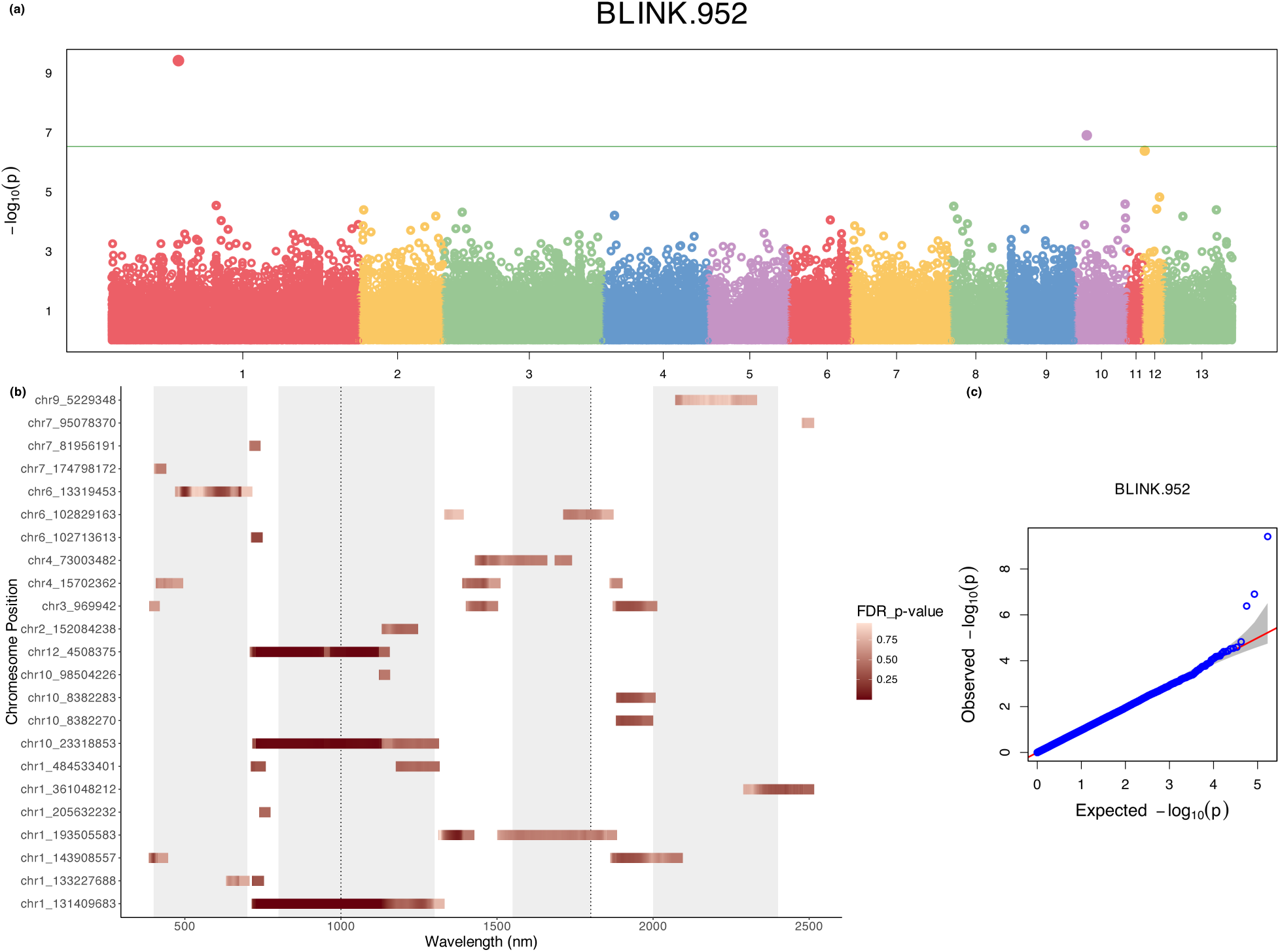
SW-GWAS results. (a) Manhattan plot showing the −*log*_10_(*p*_*value*) of SNP associations with BLINK for wavelength 952 with the strongest association. (b) Visualization of associations that have p-values less than 1 × 10^−5^. The x-axis represents the spectral wavelengths from 400 to 2500 nm, while the y-axis displays the corresponding associated SNPs. The gradient red color scale represents the FDR_adjusted p-values. (c) QQ plot of the BLINK model contrasting the observed versus the expected −*log*_10_(*p*_*value*) for the GWAS analysis on the wavelength 952 nm. The diagonal line represents the expected distribution under the null hypothesis.

Furthermore, correlations with the same SNP spanning more distant spectral regions may also reflect underlying biological processes and warrant further investigation. The associations span nearly the entire spectral range and are distributed across most chromosomes. The significance level, adjusted by *M*_*eff*_, stands at 5.0 × 10^−5^. Notably, no SNP met this threshold.

In the HSC-PA-GWAS, Figure 6a shows the associations that surpassed our exploratory threshold of 1 × 10^−5^. A total of 20 SNP associations distributed over 20 segments emerged from the analysis, all of which were also found in the SW-GWAS. Focusing on segment associations obscures spectral information, and so Figure 6b maps these associations back to their corresponding spectral wavelengths for better interpretation. This visualization facilitates the identification of specific spectral regions linked to each SNP. It is noteworthy that these associations span a range of chromosomal regions, indicating a complex genetic basis of the spectral traits. The significance level, after adjusting for *M*_*eff*_, is set at 4.7 × 10^−3^. Two associations, the SNP chr1_131409683 with Segments 37 and 38 (734-1143 nm), surpassed this threshold. The QQ plots for Segments 37 and 38, as depicted in Figure 6c and d, serve a dual purpose of validation and diagnostic assessment. Figure 6e and f show the phenotypic distributions of the significant marker (chr1_131409683) affiliated with Segments 37 and 38. In this distribution, the numbers 0, 1, and 2 represent homozygous alleles as the reference, heterozygous alleles, and homozygous alternative alleles, respectively. The majority of the RILs are assigned reference or heterozygous alleles at this site, as evidenced by the greater number of points in classes 0 and 1. A clear difference in the mean values of these three classes is evident in the boxplot.

**Figure 6.**
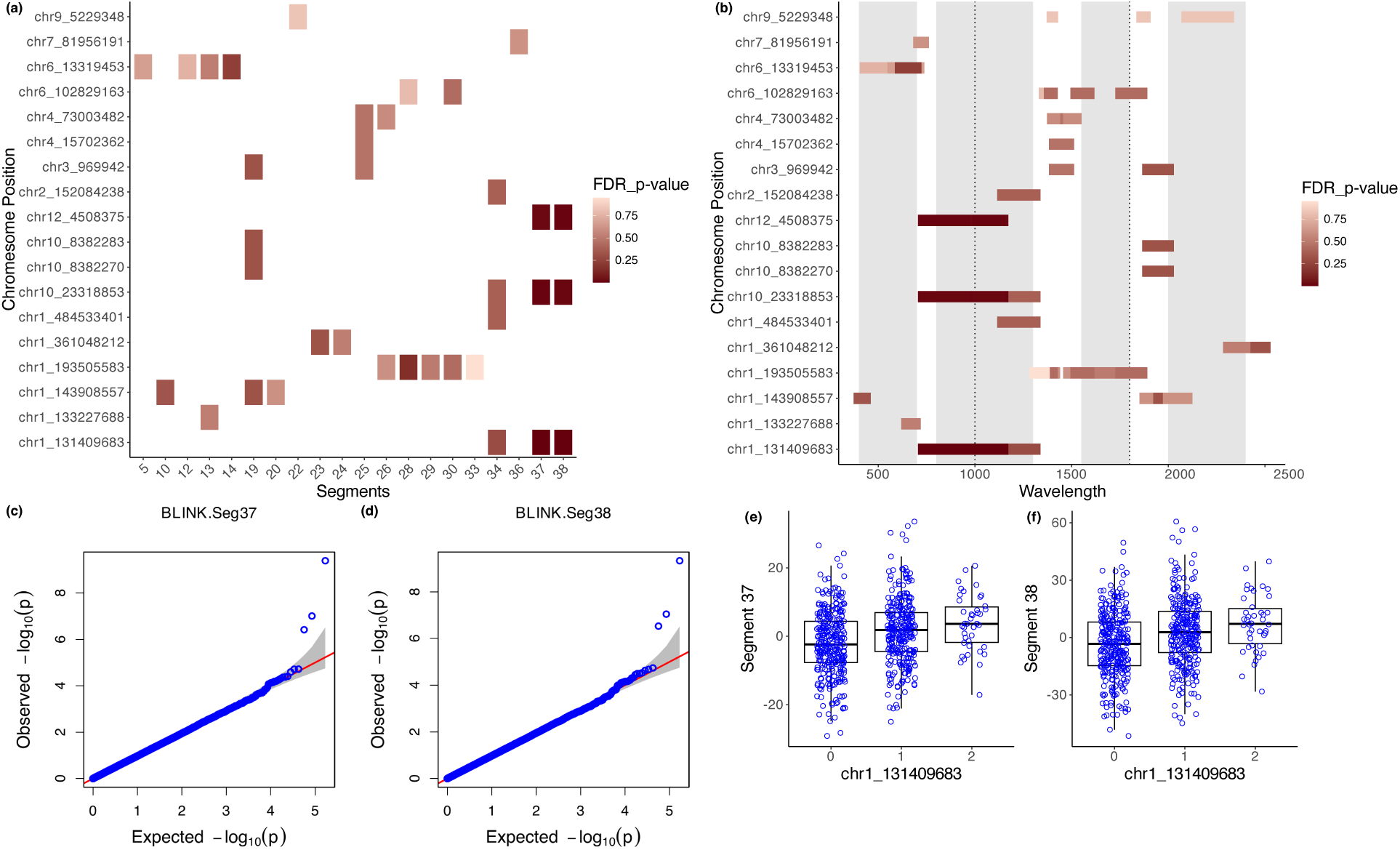
HSC-PA-GWAS results. (a) Detailed visualization of associations that surpass the exploratory threshold. The X-axis enumerates the 20 segments, while the Y-axis displays the corresponding associated SNPs. (b) An alternative representation of the associations from (a) with the X-axis showcasing the spectral wavelengths, providing insights into the specific spectral regions of the associations. (c)-(d) QQ plots highlighting the observed versus expected −*log*_10_(*p*_*value*) for GWAS analysis on Segment 37(c) or Segment 38 (d). (e)-(f) Phenotypic distribution for the significantly associated marker (chr1_131409683) in association with Segment 37(e) or Segment 38(f). 0 represents a homozygous allele from the reference, 1 a heterozygous allele, and 2 a homozygous alternative allele.

To further elucidate the genetic association indicated by HSC-PA-GWAS, we searched for potential candidate genes within a ± 100 kb window surrounding the SNP chr1_131409683. This led to the identification of one gene of interest: Niat3g_06527 (more annotation information can be found in Appendix S1: Table S2). This gene is annotated as Carbonic anhydrase (CA).

We broadened our analysis to include the top 10 SNPs from HSC-PA GWAS with the smallest FDR-adjusted P-values (see Appendix S1: Table S2). The top six associations are linked to Segment 37 or Segment 38 in the red-edge and near-infrared regions. In addition to the significantly associated locus (chr1_131409683), these associations involve two SNPs: chr12_4508375 and chr10_23318853. Furthermore, the significantly associated locus (chr1_131409683) is also associated with Segment 34 (1144–1310 nm), which lies immediately after Segments 37 and 38 and remains within the NIR region.

## 4 DISCUSSION

### 4.1 HSC-PA-GWAS for discovering genetic associations with spectral variation

Understanding the genetic foundation of leaf spectral diversity will facilitate advances in biodiversity monitoring and phenotyping, but determining appropriate approaches to extract genetic information from complex, context-dependent spectral phenotypes is a critical step (Li et al., 2023). SW-GWAS, while straightforward, is limited by the high dimensionality of spectral data and the spectral resolution of instruments, typically ranging from 3-10 nm. Subsampling can simplify data and enhance signal-to-noise ratios, yet risks omitting key data points, and the choice of bands for subsampling and the resulting number of traits require consideration and testing. It is common to subsample the continuous spectral data to 1 nm intervals, as done here for the SW-GWAS and as a basis for other analyses, because 1 nm is the subsampling interval provided in the calibrated output data from the ASD FieldSpec and other field spectrometers.

Cavender-Bares and colleagues (Cavender-Bares et al., 2016) subsampled leaf spectra at a larger interval of 5 nm prior to PLS-DA and evolutionary analysis and stated that the decision regarding the interval width (1, 2, or 10 nm) did not significantly impact the results. Czyż and colleagues used a coarser subsampling interval of 10 nm to estimate the predictive power of spectral bands for genetic structure (Czyż et al., 2020). Dimensionality reduction methods like PCA yield complex components challenging to biological interpretation (Li et al., 2023).

Similarly, spectral indices and methods like varimax rotation have limitations due to continuous spectra and multiple influencing factors. Partial least squares regression (PLSR) is another alternative, often combined with discriminant analysis or beta regression models, but it requires orthogonal trait measurements for accurate associations and carries the risk of overfitting (Cavender-Bares et al., 2016; Wang et al., 2020; Stasinski et al., 2021).

Here, we developed HSC-PA-GWAS as a new approach that retains the information in spectra in a biologically interpretable form without losing meaningful information, and without requiring an *a priori* knowledge to target specific traits, while maintaining sufficient statistical power to identify potentially meaningful associations. This method stands out for its combination of sensitivity and statistical power when handling spectral phenotypes. Associations identified using HSC-PA-GWAS were consistent with the SW-GWAS method, as every SNP identified by HSC-PA-GWAS was also detected by SW-GWAS. However, the HSC-PA method surpasses SW-GWAS in several key aspects. One of its strengths is its ability to reduce data dimensionality while accounting for phenotypic correlations within spectra, by transforming multiple correlated wavelengths into a single value, typically represented by the first principal component. This mitigates the challenges of multiple testing and correlated phenotypic data and thus increases statistical power. By aggregating information across multiple wavelengths, the HSC-PA method offers a more comprehensive and potentially more accurate representation of the underlying traits with its ability to discover the correlation structure among wavelengths from data. The resulting genetic associations encompass a broader portion of the spectrum than those derived from indices, which focus on specific wavelength combinations, or from single wavelengths, which do not behave independently. While indices remain valuable for highlighting proxies of relevant biological processes or capturing relative changes between spectral regions, the HSC-PA approach retains a wider spectral context for interpretation. Moreover, compared to spectral subsampling (e.g., sampling or averaging every 10 nm), which typically does not allow for (varying) covariation patterns of spectral bands and focuses only on neighboring bands, the HSC-PA method provides a more data-driven approach that accounts for empirical internal correlations.

Comparing the HSC-PA method with established methods, such as an approach combining partial least squares regression (PLSR) and linear regression to interpret spectral features in terms of other traits (Verrelst et al., 2019), indicates that HSC-PA has an advantage in at least two ways: supporting human interpretation of data-derived spectral features and thus genetic associations, and allowing for experimental designs aimed at discovering associations with novel trait variation or where there are limitations on producing appropriate validation datasets required for regression. Verrelst and colleagues (Verrelst et al., 2019) also emphasized the importance of nonparametric regression and machine learning methods, like the Kernel Ridge Regression (KRR) and Gaussian Processes Regression (GPR), for their ability to capture nonlinear relationships in spectral data. These methods, while powerful, often require careful tuning, can be computationally intensive, and carry the danger of overfitting. The HSC-PA method offers a data-driven yet human-interpretable approach with the caveat that its criterion for simplification is linear (dimensionality in a PCA). As discussed in studies on human facial shapes, from which the HSC-PA method was adapted (Claes et al., 2018), this methodological shift allows for a transition from a global to a local understanding of spectral variation, and a holistic view of its genetic basis.

### 4.2 Spectra adjustment and genetic associations

In this study, we used replicated plants of a single tester genotypes of *N. attenuate* as “phytometers” to account for small-scale (2×2 planting positions) spatial variation across an experimental field. In addition, we accounted for differences in other environmental variables such as measurement time as well as for maternal effects. By applying a linear model across the full spectral range and using the residuals as adjusted spectra for downstream analyses, we were able to reduce the influence of confounding environmental variables. This correction, which could not have been incorporated directly into a mixed-model GWAS analysis, substantially altered many results, including derived phenotypes and GWAS associations, leading to different segment definitions and candidate genes in comparison to directly conducting GWAS on spectral data without prior correction for non-genetic effects (Appendix S1: Supporting Text, Results from raw spectral data). Such shifts underscore the importance of accounting for non-genetic effects in spectral-genomic studies, as failing to do so can lead to markedly different—and potentially misleading—biological conclusions. Here, we were able to do so both by using a phytometer and by accounting for environmental factors dictated by the experimental and measurement design; while a genetically invariable phytometer is rarely an option in ecological studies, it should usually be feasible to account for quantitative environmental and experimental effects on phenotype values.

Advanced genetic association models considering epistasis, akin to HSC-PA’s handling of spectral correlations, could enhance GWAS power, sensitivity, and interpretability, with advances in machine learning opening new possibilities (Libbrecht and Noble 2015; Wu et al., 2021). Random forest models have already been used for analyzing complex genetic effects on multivariate phenotypes, though their significance interpretation differs (Brieuc et al., 2018; Wang et al., 2013). The “Epi-MEIF” approach by Saha et al., (2022), using conditional inference forests for epistasis in complex phenotypes, offers a method for significant association identification, accommodating multiple loci testing as an add-on to conventional single-locus models like those in GAPIT.

4.3 Ecological significance of findings in *N. attenuata*

We discovered a significant association between the SNP chr1_131409683 and Segments 37 and 38 (734-1143 nm) in the HSC-PA-GWAS. One interesting candidate gene was identified within 100 kb of this SNP: Niat3g_06527, is annotated as a carbonic anhydrase (CA). Carbonic anhydrases catalyze the reversible hydration of CO₂ to bicarbonate and protons (CO₂ + H₂O ⇌ HCO₃⁻ + H⁺) (Supuran, 2008). In plants, multiple CA isoforms are found in the chloroplast stroma, thylakoid membranes, cytosol, and other compartments (Fabre et al., 2007; Rudenko et al., 2015). Chloroplastic CAs are thought to facilitate the rapid supply of CO₂ to Rubisco by accelerating the interconversion between dissolved inorganic carbon species, thereby supporting the Calvin–Benson cycle (Badger & Price, 1994). In C₄ and CAM plants, CA activity in mesophyll cells is also critical for providing bicarbonate to phosphoenolpyruvate carboxylase during the initial fixation step (Hatch & Burnell, 1990). In C₃ species such as *N. attenuata*, stromal and thylakoid-associated CAs have additionally been implicated in maintaining photosynthetic efficiency under fluctuating light and CO₂ conditions, and in the regulation of pH and ion balance in the thylakoid lumen (Tiwari et al., 2016).

The associated spectral range, 734-1143 nm, spans the red-edge and near-infrared regions, which are sensitive to changes in leaf internal structure and chlorophyll content. Genetic variation in a chloroplast-localized CA could influence CO₂ assimilation rates and downstream effects on chlorophyll concentration, mesophyll cell structure, or water status, any of which can alter reflectance in this part of the spectrum. The link between Niat3g_06527 and these spectral regions therefore suggests that natural allelic differences in CA function may contribute to variation in photosynthetic performance and leaf optical properties in *N. attenuata*.

In the context of the ecology of *N. attenuata*, understanding the genetic basis of spectral variation can provide insights into the adaptive mechanisms of this wild plant. The significant association of a SNP close to a gene annotated as carbonic anhydrase, with a spectral range spanning the red-edge and near-infrared regions—wavelengths closely linked to chlorophyll content, canopy structure, and photosynthetic capacity—may reflect evolutionary pressures that have shaped the species’ genetic architecture to optimize carbon fixation and water-use efficiency, enhancing its survival in the variable and often harsh environments of its native habitat. Our extended analysis of the top 10 SNPs has unveiled additional genetic associations that may play important significant roles in the ecological adaptation and survival of *N. attenuata*.

Two other statistically significant associations were discovered with the *ARDSI*_*C*_w_index, an index of water content that uses the wavelengths 1080, 1240, 1360, and 1560 nm, which are mostly (except for 1080 nm) in the range of wavelengths longer, *i.e.*, further into the infrared region, than those comprising segments 37 and 38. The two associated SNPs, both also on chromosome 1, were located in the vicinity of the gene Niat3g_08138 (chr1_193505583), annotated as a Zeta toxin domain-containing protein, and the genes Niat3g_17277 annotated as TPR_REGION domain-containing protein and Niat3g_17282 annotated as Exocyst subunit Exo70 family protein (chr1_438155938). Proteins with these domains are documented to serve a wide variety of functions in defense, growth and development, and cellular function.

It is interesting that the strongest associations were all identified between loci on chromosome 1 and wavelengths longer than those of visible light, in the red edge to infrared portions of the leaf spectrum, that are influenced by micromorphology, structure, abundant non-pigment organic constituents, and water content. The strongest associations were within regions of high signal:noise in spectral reflectance, *i.e.*, where the detection of phenotypic differences should be most robust.

### 4.4 Limitations and outlook

One limitation of our study lies in the phenotypic interpretation for biological meanings specific to the samples we used. The six spectral indices we selected are derived from existing literature and have not been validated in *N. attenuata* generally, or for this dataset. Without chemical or other analyses to specifically quantify the constituents represented in the indices, such as chlorophyll, our interpretations remain speculative. Establishing causal links between candidate genes and specific biological processes is inherently difficult, particularly in ecological systems where multiple interacting factors may influence spectral traits (Hoban et al. 2016). Moreover, while the HSC-PA-GWAS approach has identified potential candidate genes, any influence of these genes on the traits of interest remains to be verified, e.g. through the generation and testing of knock-out, knock-down, and overexpression lines.

While GWAS with a MAGIC population does not require replication of genotypes, the absence of such replicates prevented us from testing variation between genotypes directly using the analysis of variance approach. Nevertheless, by removing small-scale environmental variation using replicated phytometers of a single genotype as well as other environmental variation and maternal effects, the residual phenotypic differences between plants was likely due to their genetic differences rather than further environmental causes.

Our study noted unexpectedly high heterozygosity levels (mean around 0.5, see Appendix S1: Figure S3) in the dataset, potentially due to the low-pass sequencing (mean 0.5x coverage) of the RILs (Ray et al., 2023). The significant data gaps (about 95% missing data) necessitated imputation from parental lines, which weren’t highly inbred, likely inflating heterozygosity estimates in the RILs. While the imputed genotype matrix was accurate for known QTLs (Ray et al., 2023), the sequencing and imputation approach introduces uncertainty in true heterozygosity levels. Factors like residual heterozygosity and heterozygote hotspots, seen in species like maize (Liu et al., 2018), may also influence these high heterozygosity readings in the *N. attenuata* MAGIC RIL population. Future studies with higher sequencing depth could more accurately determine homozygosity and heterozygosity in this population, aiding in dissecting genetic architecture and validating GWAS findings.

Looking forward, there is great potential for deepening our understanding of leaf spectral data using the *N. attenuata* system. Our analysis indicates relatively large variance in blue light reflectance and perhaps related photosynthetic traits among the wild genotypes contributing to the *N. attenuata* MAGIC population, as discussed previously (Li et al., 2023). Variation in conserved and essential traits, which can be mediated by variants of regulatory genes embedded in coexpression networks (Ray et al., 2023), may lend different genotypes a competitive advantage under changing environmental conditions. The significant associations identified here thus indicate genotype and phenotype targets for novel and interesting in-depth functional studies that could help us to better understand plant survival strategies. A combination of natural variants, genetic modification, and laboratory assays, such as gene expression analyses (Schena et al., 1995), protein-protein interactions (Fields and Song 1989), and chromatin immunoprecipitation assays (Solomon et al., 1988), comprise the gold standard for functional testing and validation of discovered genetic associations. At this stage, the associations reported here represent hypotheses that should be subjected to these tests.

Unraveling the genetic basis of leaf spectral properties has far-reaching implications and could for example revolutionize plant biodiversity monitoring, ecophysiology, and adaptation research, and help to identify genotypes resilient to stresses associated with global change. Our study sets the stage for future research aimed at deciphering these links by presenting a widely applicable new method for data-driven discovery of associations which can be integrated with existing and novel tools to discover genetic associations and support moving from association to prediction or causation.

## 5 CONCLUSIONS

In our study, we explored the genetic basis of leaf spectral properties in *N. attenuata* by associating spectral phenotypes with SNP genotypes in GWAS during a field experiment. We analyzed leaf reflectance spectra from 400 to 2500 nm, identifying regions correlating with leaf composition and function. We developed the Hierarchical Spectral Clustering with Parallel Analysis (HSC-PA) method to manage the data’s high dimensionality and correlation, proving more effective than other methods like spectral indices or individual wavelengths. This approach offered greater statistical power and maintained interpretable features, despite reduced dimensionality.

Our findings included a significant SNP association in the red-edge to near-infrared spectral range (734–1143 nm), a region sensitive to chlorophyll content, canopy structure, and photosynthetic efficiency. The most significant SNP lies near the Niat3g_06527 gene, annotated as carbonic anhydrase, an enzyme central to photosynthetic carbon fixation and stomatal regulation. These results require further validation, but they raise the intriguing possibility that variation in red-edge/NIR reflectance could be linked to differences in carbon assimilation and water-use strategies in *N. attenuata*, potentially reflecting adaptations to its native, resource-variable environments.

This study introduces a method and framework that can be broadly applied to attain deeper understanding of the genetic underpinnings of leaf spectral properties. The associations identified here using an advanced genetic toolset built for a native plant, and studied in a field experiment within the plant’s natural range, have the potential to advance our understanding of the genetic mechanisms underlying photosynthetic efficacy and related growth and defense strategies in plants, and could support new insights in the application of remote sensing to biodiversity assessment.

## ACKNOWLEDGMENTS

The authors acknowledge funding from the NOMIS Foundation and the University of Zurich, including the University Research Priority Program on Global Change and Biodiversity, and the Max Planck Society. We thank our division in the Department of Geography at the University of Zurich, the Remote Sensing Laboratories, for support and for the shared use of equipment. We thank the local managers at WCCER, N. Carlson and R. Carlson, and the 2019 field crews for plant cultivation and plot management. We thank the WCCER for supporting these experiments.

## 6 AUTHOR’S CONTRIBUTIONS

CRediT taxonomy roles are listed with authors in alphabetical order. Conceptualization: I.T.B., M.C.S., M.E.S. Data curation: B.S., C.L., R.R. Formal analysis: B.S., C.L. Funding acquisition: I.T.B., M.E.S. Investigation: E.A.C., M.C.S. Methodology: B.S., C.L., E.A.C., M.C.S. Project administration: M.C.S. Resources: I.T.B., M.E.S., R.H. Supervision: M.C.S. Visualization: C.L. Writing—original draft: C.L. Writing—review and editing: B.S., C.L., E.A.C., I.T.B., M.C.S., M.E.S., R.H., R.R.

## 7 CONFLICT OF INTEREST STATEMENT

The authors declare no conflicts of interest.

## Notes

### Competing Interest Statement

The authors have declared no competing interest.

### Summary of Updates

This version of the manuscript has been substantially revised following editorial and reviewer feedback to improve clarity, readability, and compliance while maintaining its focus on developing and applying methodological approaches to link plant genetic variation with leaf spectral traits in Nicotiana attenuata. A major adjustment introduced during the revision was the use of reference genotypes, grown repeatedly across the field (one in every four planting positions), as controls to account for environmental effects such as spatial position, measurement time, and maternal cytoplasmic effects. We applied a linear model across the full spectral range and used the residuals as adjusted spectra for downstream analyses. This change substantially affected the results, including phenotypes and GWAS associations, and led to different segment definitions and candidate genes.

