## Appendix for "Genome-wide associations of leaf spectral variation in MAGIC lines of *Nicotiana attenuata*"

Manuscript title: Association of leaf spectral variation with functional genetic variants

### Supporting Text

#### Plant Material and Field Site

*Nicotiana attenuata* is a native tobacco species predominantly found in the southwestern United States. The field site for this research was located at the Walnut Creek Center for Education and Research (WCCER) in Prescott, Arizona, within the natural habitat of *N. attenuata*. The MAGIC population of *N. attenuata* was derived from the Utah (UT) accession, a 31$^{st}$-generation inbred line originally collected from the Desert Inn ranch in Washington County, Utah, USA. Other natural accessions used in this study were detailed in a previous publication (Ray et al., 2023). Germination and cultivation for the Arizona field plantation were previously described in (Li et al., 2023).

#### Generation of MAGIC RIL Population, DNA sequencing, and genome annotation

To structure the genetic diversity in the MAGIC RIL population, a crossing scheme was developed with 26 parental lines. This entailed five rounds of systematic inter-crosses, ensuring that each offspring would have all 26 parental lines as ancestors. The first round involved diallelic crossing of each of the 26 parental lines with each other, resulting in an F1 or A- generation of 325 different crosses. This process was detailed, with specific procedures to prevent self-pollination and ensure accurate cross-pollination. The subsequent rounds (B- to E-generations) further diversified the genetic makeup of the offspring. By the end of the fifth round, each plant had all 26 MAGIC founders as parents. These systematic crosses were followed by six generations of inbreeding to produce Recombinant Inbred Lines (RILs), resulting in a MAGIC RIL population consisting of two replicates of 325 RILs. For the sequencing of the 650 MAGIC RILs, seedlings from the L1 and L2 generations were grown in the GH, and the sequencing was conducted at Novogene HK (Ray et al., 2023; Li et al., 2023).

To associate the resulting phenotypic data with genetic variation, we perform genotype calling from low-coverage sequencing data of the RILs. This involves using the GATK HaplotypeCaller for individual variant discovery, followed by the aggregation of these variants using GATK CombineGVCFs (DePristo et al., 2011). Subsequent filtering and processing yield a matrix of SNPs, which are then imputed to account for missing data, resulting in a pool of 183,942 SNPs. We further removed SNPs that showed no variance, eliminating 15,039 SNPs and leaving 168,903 SNPs for downstream analyses.
Genome annotation was performed using an RNA-Seq–guided approach. Publicly available RNA-Seq data were aligned to the genome, and transcript models were assembled and refined using StringTie and Mikado, incorporating splice junction information from Portcullis. These transcriptomes, together with curated protein evidence from SwissProt, were integrated into the MAKER annotation pipeline to iteratively improve and predict gene models. Functional annotations were assigned through homology searches against TAIR11, SwissProt, and TrEMBL using DIAMOND, and gene descriptions and GO terms were generated with the AHRD pipeline.

### Equipment and optical measurement

We employed a FieldSpec 4 spectroradiometer (Analytical Spectral Devices, Inc., Malvern Panalytical) 18140 with a plant probe (S/N 445) and leaf clip V2 attachment for measuring leaf optical properties. The FieldSpec encompasses three detectors, covering the visible and near-infrared to the shortwave infrared range of electromagnetic radiation. The devices offer a spectral resolution of 3 nm at 700 nm and 10 nm at 1400 and 2100 nm, and the FieldSpec system is radiometrically calibrated to provide measurements from which values for radiance and subsequently reflectance can be derived for every nanometer between 350 and 2500 nm.

Mature, hydrated, cut leaves harvested from comparable positions of field-grown plants were measured in batches immediately after each harvest, at their widest part, avoiding the midvein. To completely cover the field of the leaf clip, either one or two leaves were utilized, as described in (Li et al., 2023). Each sample underwent 20 scans under four conditions: white background reference (WR), white background with leaf (WRL), black background reference (BR), and black background with leaf (BRL). All measurement procedures for the Arizona field study were previously described (Li et al., 2023).

### Data processing

Data processing was executed using R (R Core Team 2023), primarily employing the spectrolab package (version 0.0.10) (Meireles et al., 2017). We employed a rigorous three-step filtering approach to remove outliers from our dataset, as previously described (Li et al., 2023). Briefly, we initially conducted a visual inspection of different measurement types, followed by the application of the Local Outlier Factor (LOF) method for each type. A final visual check was performed after calculating reflectance, ensuring the exclusion of outliers that could strongly impact our analysis. The spectral range analyzed spanned from 400-2500 nm, excluding the initial 50 nm (350-400 nm) due to high measurement uncertainty (Petibon et al., 2021). The calculated reflectance (CR) of a sample was obtained from the mean of scans using the following formula from (Miller et al., 1992):

$$CR = (R_{WR} \cdot R_{BRL} - R_{BR} \cdot R_{WRL})/(R_{WR} - R_{BR})$$

### Genome-wide association studies (GWAS)

We applied three approaches—Index-GWAS, Single Wavelength GWAS (SW-GWAS), and Hierarchical Spectral Clustering with Parallel Analysis GWAS (HSC-PA GWAS)—to the spectral dataset (616 individuals × 2101 spectral phenotypes). As described in the main text, the raw spectral data were adjusted using a linear model. For comparison, we also performed the same GWAS analyses on the unadjusted raw spectra, with selected results presented in this Appendix: Supporting text: Results from raw spectral data.

### Index-GWAS

This approach uses spectra to simultaneously assess multiple phenotypes defined by six common spectral indices that are widely recognized for their ability to characterize various vegetation traits. These indices were chosen based on their sensitivity to specific vegetation properties and their proven applicability in previous research (Wan et al., 2021), as summarized in Table 1:

The background and significance of these indices are further elaborated below:

1. Normalized Difference Water Index ($NDWI$): $NDWI$ is used to assess vegetation water content. It is based on the differential absorption of water in the near-infrared and shortwave infrared regions of the spectrum. $NDWI$ has been widely applied in remote sensing to detect water stress in plants (Gao 1996; Zhang and Zhou 2019).
2. Red-edge chlorophyll index ($CI_{re}$): $CI_{re}$ is designed to estimate vegetation chlorophyll content. The red-edge region of the spectrum is sensitive to chlorophyll concentration, making $CI_{re}$ a valuable index for assessing plant vigor and growth (Gitelson et al., 2001; Qian et al., 2021).
3. Chlorophyll Carotenoid Index ($CCI$): $CCI$ is sensitive to carotenoid/chlorophyll ratio and was initially used to track photosynthetic phenology (Gamon et al., 2016).

We additionally used three adjusted ratio of difference spectral indices ($ARDSI$), which are designed to reduce the differences in estimates obtained for adaxial and abaxial leaf surfaces (Wan et al., 2021), for assessing

1. vegetation chlorophyll content ($C_{ab}$),

2. water content ($C_w$), and
3. dry matter content ($C_m$), which have been validated in other plant species at both leaf and canopy levels.

### Data-driven approaches

The data-driven approach employed two different methods to identify genetic variants associated with leaf reflectance spectra (ranging from 400 to 2500 nm) in the RILs:

1. Single Wavelength GWAS (SW-GWAS): This approach treats each wavelength as a separate phenotype and runs a GWAS on each wavelength. This allows for the identification of genetic variants associated with specific wavelengths of the leaf reflectance spectra. We note that the resolution of the spectrometer is reported by Analytical Spectral Devices (ASD, Inc.) to range from 3-10 nm with the narrowest resolution for the shortest wavelengths, and so neighboring wavelengths may not behave independently.
2. Hierarchical Spectral Clustering with Parallel Analysis GWAS (HSC-PA-GWAS): This approach, adapted from a method for GWAS on human facial shapes (Claes et al., 2018; Sero et al., 2019), reduces the dimensionality of the data by clustering the spectra into segments based on their co-behavior. The first principal component (PC) of each segment, which captures the majority of the variation within the segment, was then used as a phenotype for GWAS. The number of segments and the number of PCs to retain for each segment were determined using Parallel Analysis (PA), a statistical method that compares the observed eigenvalues with those obtained from random data (Franklin et al., 1995; Hayton et al., 2004).

### Software and setting

The Linear models were build in R using the base package lm() function.

A series of GWA studies were conducted in R using the Genome Association and Prediction Integrated Tool (GAPIT) version 3 (Wang and Zhang 2021) using the following models:

1. Generalized Linear Model (GLM) (Price et al., 2006): A model that captures the linear relationship between genetic markers and phenotypic traits using a flexible generalization of ordinary linear regression that allows for response variables that have error distribution models other than a normal distribution. GLM does not incorporate kinship.
2. Mixed Linear Model (MLM) (Yu et al., 2006): Extends the GLM by incorporating both fixed effects of population structure and random effects of kinship to control for spurious associations, which, however, also reduces power to detect true associations which may e confounded with population characteristics.
3. Fixed and random model Circulating Probability Unification (FarmCPU) (Liu et al., 2016): Iteratively fits fixed and random effects to improve the power and precision of GWAS. FarmCPU eliminates the confounding effect of kinship by utilizing a fixed-effect model. The kinship derived from the associated markers is then used to select the associated markers using the maximum likelihood method. This method effectively overcomes the problems of model overfitting that arise with stepwise regression.
4. Bayesian information and Linkage-disequilibrium Iteratively Nested Keyway (BLINK) (Huang et al., 2019): Iteratively incorporates associated markers as covariates to eliminate their connection to the cryptic relationship among individuals. The associated markers are selected according to linkage disequilibrium, optimized for Bayesian information content, and reexamined across multiple tests to reduce false negatives (Wang and Zhang 2021).

Testing multiple methods rather than choosing one helps to assess our association procedures. Different models have varying strengths depending on the nature of the data and the underlying genetic architecture of the trait being studied. By employing a suite of models, we aimed to cross-validate results and potentially uncover associations that might be missed by a single model.

### Heritability estimation

Narrow-sense heritability ($h^2$) of each trait was estimated using the variance components output by the Mixed Linear Model (MLM) in GAPIT (Zhang et al., 2010), providing a measure of the genetic contribution to the observed phenotypic variance. GAPIT partitions the total phenotypic variance into additive genetic variance ($\sigma_g^2$) and residual variance ($\sigma_e^2$), and calculates heritability as:

$$\mathrm{h}^2 = \frac{\sigma_g^2}{\sigma_g^2 + \sigma_e^2}$$

Estimates were obtained from the variance components reported in the “Optimum_MLM” output files.

The estimated heritability across most phenotypes was 0. Among the six indices, $CCI$ showed the highest value at 15.3%, followed by $ARDSI\_C_m$ (6.3%) and $ARDSI\_C_{ab}$ (5.8%), while $NDWI$, $CI_{re}$, and $ARDSI\_C_w$ were estimated at 0. For the single-wavelength GWAS, heritability was 0 for most wavelengths, with only 27 wavelengths (402, 405, 2469–2470, 2475–2477, and 2481–2500 nm) showing non-zero values ranging from 1% to 1.7%. For the HSC-PA approach, heritability across all spectral segments was estimated at 0. These results may reflect limitations inherent to our dataset. First, the MAGIC RIL population was sequenced at very low coverage (~0.5×), which introduced a high proportion of missing data (~95%) that had to be imputed from parental genotypes. Since the parental lines themselves were not highly inbred, this imputation likely inflated heterozygosity estimates and reduced the accuracy of SNP calls. As a consequence, the genetic relationship matrix used to partition variance may not fully capture the true additive genetic variance. In addition, the experimental design of the RILs, with each line represented by a single plant rather than replicated individuals, further constrains the ability to disentangle genetic from residual variance. Taken together, these factors likely explain why heritability estimates were close to zero for most traits and should be interpreted with caution.

### Results from raw spectral data

The distribution of the six spectral indices calculated from the raw spectra is shown in Figure S1a. Figures S1b and S1c present the results from HSC-PA. The raw spectra were segmented into 40 distinct patterns, of which 21 segments each retained one principal component, collectively spanning the entire spectrum.

The effective number of independent traits, calculated using the PhenoSpD method, was 5.0 for the indices, 11.8 for HSC-PA, and 1042.2 for the single wavelengths.

Table S3 shows the top 10 significant SNPs with candidate gene annotations from the HSC-PA-GWAS on raw spectral data. Notably, two closely located SNPs, chr10_8382283 and chr10_8382270, were significantly associated with Segment 19 and Segment 22, respectively.

Jasmonate Signaling Is Embedded in an Herbivory-Elicited Gene Network.” Proceedings of the National Academy of Sciences 120 (35): e2308500120.

Wang, Jiabo, and Zhiwu Zhang. 2021. “GAPIT Version 3: Boosting Power and Accuracy for Genomic Association and Prediction.” Genomics, Proteomics & Bioinformatics 19 (4): 629–40.

Table S1. Spectral Ranges of All Segments

| Segment | Range | Segment | Range |
|---|---|---|---|
| 1 | 400-709, 1359-2500 | 20 | 1880-1893, 2001-2096 |
| 2 | 710-1358 | 21 | 2312-2500 |
| 3 | 400-709, 1400-1522, 1864-2500 | 22 | 1400-1401, 1864-1879, 2097-2311 |
| 4 | 1359-1399, 1523-1863 | 23 | 2397-2500 |
| 5 | 518-614, 695-709 | 24 | 2312-2396 |
| 6 | 400-517, 615-694, 1400-1522, 1864-2500 | 25 | 1412-1484 |
| 7 | 400-517, 615-694 | 26 | 1402-1411, 1485-1522 |
| 8 | 1400-1522, 1864-2500 | 27 | 1381-1399, 1523-1863 |
| 9 | 433-517, 615-694 | 28 | 1359-1380 |
| 10 | 400-432 | 29 | 1381-1385, 1590-1754 |
| 11 | 615-694 | 30 | 1386-1399, 1523-1589, 1755-1863 |
| 12 | 433-517 | 31 | 1144-1358 |
| 13 | 648-690 | 32 | 710-1143 |
| 14 | 615-647, 691-694 | 33 | 1311-1358 |
| 15 | 1400-1401, 1864-2500 | 20 | 1880-1893, 2001-2096 |
| 16 | 1402-1522 | 21 | 2312-2500 |
| 17 | 1880-2096 | 22 | 1400-1401, 1864-1879, 2097-2311 |
| 18 | 1400-1401, 1864-1879, 2097-2500 | 23 | 2397-2500 |

Table S2. Top Ten Significant SNPs with Candidate Gene Annotations (HSC-PA, adjusted spectra)

| SNP | Segment | Spectral range | p_value | FDR_Adjusted_p_value | GeneID | Description | Strand |
|---|---|---|---|---|---|---|---|
| chr1_131409683 | 37 | 734-744, 946-1014, 1129-1143 | 4.07E-10 | 6.88E-05 | Niat3g_06527 | Carbonic anhydrase | + |
| chr1_131409683 | 38 | 745-945, 1015-1128 | 4.65E-10 | 7.86E-05 | Niat3g_06527 | Carbonic anhydrase | + |
| chr1_131409683 | 34 | 1144-1310 | 1.74E-06 | 2.93E-01 | Niat3g_06527 | Carbonic anhydrase | + |
| chr1_193505583 | 28 | 1359-1380 | 9.34E-07 | 1.58E-01 | Niat3g_08137 | Unknown protein | + |
| chr1_193505583 | 28 | 1359-1380 | 9.34E-07 | 1.58E-01 | Niat3g_08138 | Zeta_toxin domain-containing protein | + |
| chr10_23318853 | 37 | 734-744, 946-1014, 1129-1143 | 9.85E-08 | 8.32E-03 | Niat3g_78151 | ATP-dependent DNA helicase | - |
| chr10_23318853 | 38 | 745-945, 1015-1128 | 2.95E-07 | 1.66E-02 | Niat3g_78151 | ATP-dependent DNA helicase | - |
| chr10_23318853 | 37 | 734-744, 946-1014, 1129-1143 | 9.85E-08 | 8.32E-03 | Niat3g_78161 | 2-isopropylmalate synthase A-like | - |
| chr10_23318853 | 38 | 745-945, 1015-1128 | 2.95E-07 | 1.66E-02 | Niat3g_78161 | 2-isopropylmalate synthase A-like | - |
| chr12_4508375 | 38 | 745-945, 1015-1128 | 9.02E-08 | 7.62E-03 | Niat3g_81493 | Unknown protein | - |
| chr12_4508375 | 37 | 734-744, 946-1014, 1129-1143 | 3.84E-07 | 2.16E-02 | Niat3g_81493 | Unknown protein | - |
| chr6_13319453 | 14 | 615-647, 691-694 | 1.39E-06 | 2.35E-01 | Niat3g_70143 | GP-PDE domain-containing protein | - |

Table S3. Top 10 Significant SNPs with Candidate Gene Annotations (HSC-PA, adjusted spectra)

| SNP | Segment | Spectral range | p_value | FDR_Adjusted_p_value | GeneID | Description | Strand |
|---|---|---|---|---|---|---|---|
| chr10_8382283 | 19 | 1894-1999 | 1.81E-09 | 3.06E-04 | Niat3g_77674 | Protein quirky | + |
| chr10_8382270 | 22 | 400-422 | 1.32E-08 | 2.24E-03 | Niat3g_77674 | Protein quirky | + |
| chr3_177473323 | 19 | 1894-1999 | 1.46E-07 | 1.23E-02 | Niat3g_26415 | C2 domain-containing protein | - |
| chr3_177473323 | 19 | 1894-1999 | 1.46E-07 | 1.23E-02 | Niat3g_26416 | C2 domain-containing protein | + |
| chr3_177473323 | 19 | 1894-1999 | 1.46E-07 | 1.23E-02 | Niat3g_26417 | Bag family molecular chaperone regulator 6 | - |
| chr3_177473323 | 19 | 1894-1999 | 1.46E-07 | 1.23E-02 | Niat3g_26418 | rRNA-processing protein FCF1 homolog | + |
| chr3_177473323 | 19 | 1894-1999 | 1.46E-07 | 1.23E-02 | Niat3g_26419 | Glycos_transf_1 domain-containing protein | - |
| chr6_13319453 | 30 | 504-519 | 3.30E-07 | 5.57E-02 | Niat3g_70143 | GP-PDE domain-containing protein | - |
| chr6_13319453 | 29 | 445-503, 659-687 | 1.15E-06 | 1.05E-01 | Niat3g_70143 | GP-PDE domain-containing protein | - |
| chr6_13319453 | 28 | 610-658, 688-695 | 6.74E-07 | 1.14E-01 | Niat3g_70143 | GP-PDE domain-containing protein | - |
| chr8_96230462 | 21 | 423-444 | 2.04E-06 | 1.73E-01 | Niat3g_77194 | Integrase_H2C2 domain-containing protein | + |
| chr8_96230462 | 21 | 423-444 | 2.04E-06 | 1.73E-01 | Niat3g_77195 | Integrase_H2C2 domain-containing protein | + |

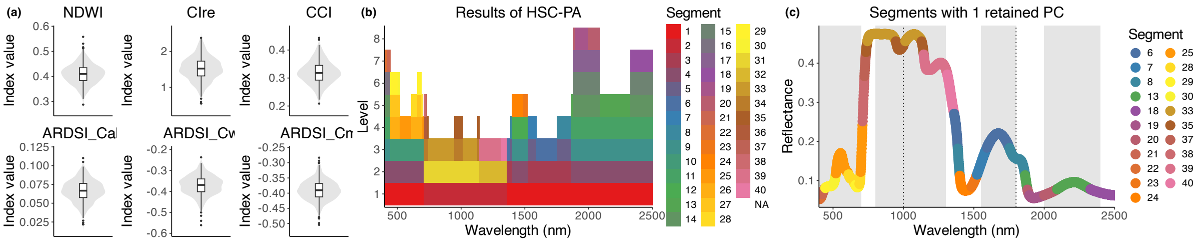


Figure S1. Raw spectra as phenotypes. To compare with Figure 2. (a) Distribution of the six spectral indices ustilized in the Indexices-GWAS approach. (b) Results from the Hierarchical Spectral Clustering with Parallel Analysis (HSC-PA). A total of 40 segments, differentiated by color, emerged from the clustering. (c) Presentation of the 21 segments, shaped as example spectra, each maintaining one retained Principal Component (PC).

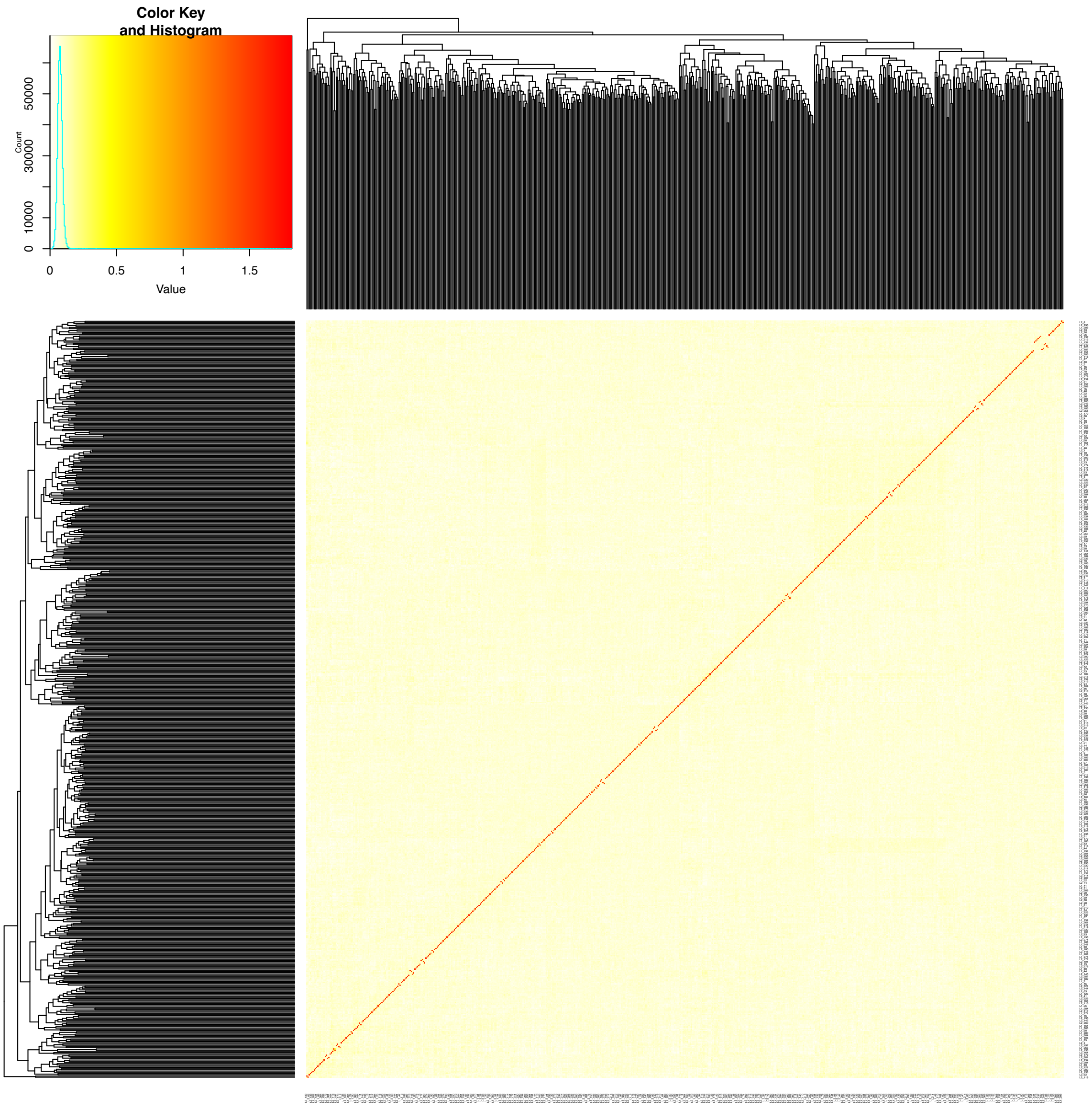


Figure S2. A heatmap of the kinship matrix indicating the relationship between individuals.

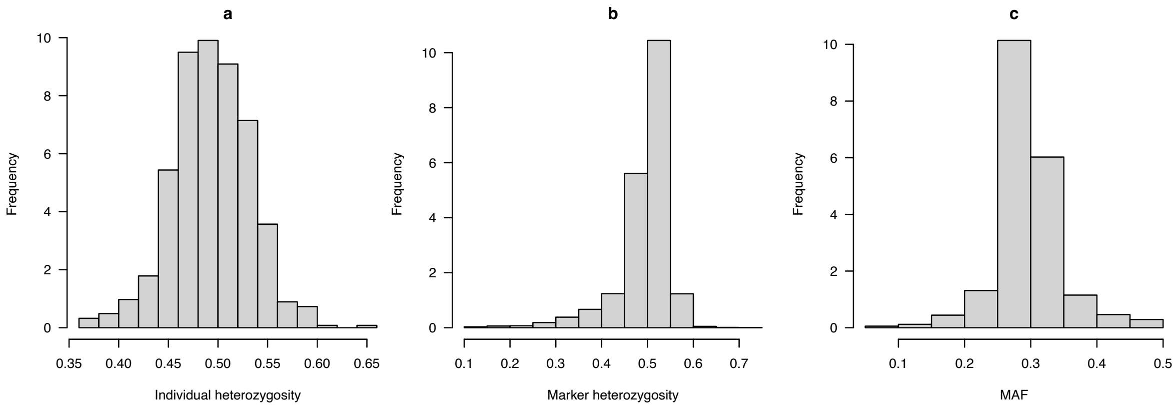


Figure S3. Frequency of heterozygosity of individual and markers.

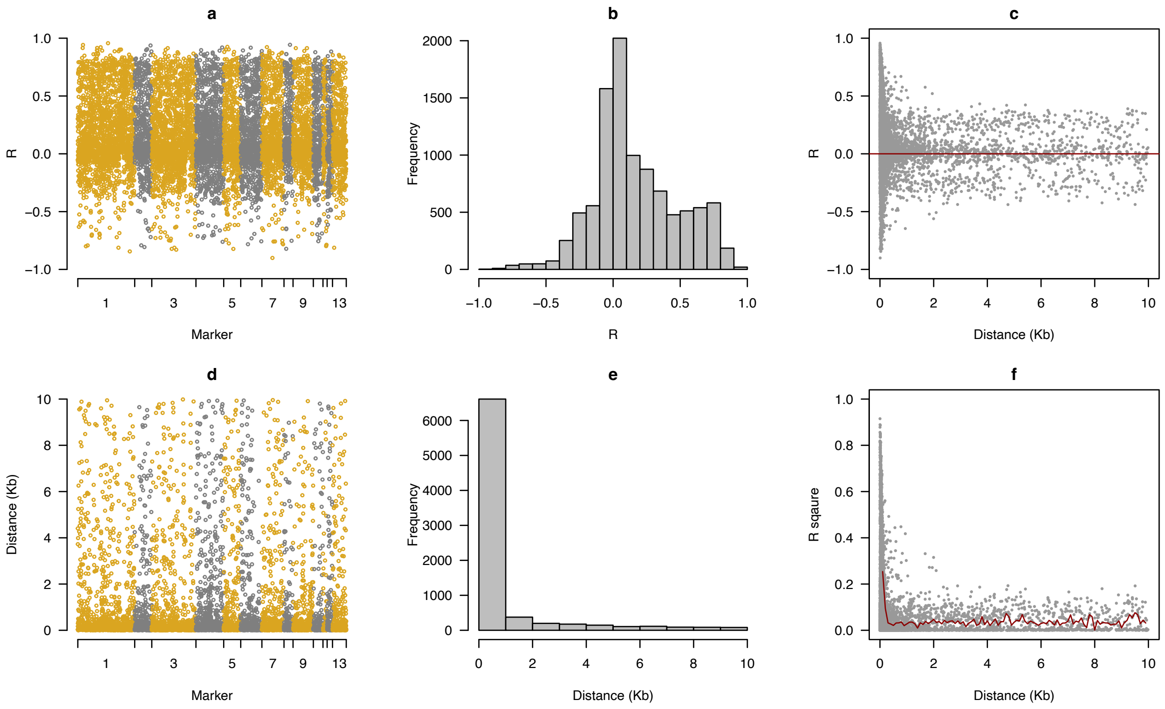


Figure S4. Distance and linkage disequilibrium (LD) decay between neighbor markers.
